# Development of a micro-CT mouse holder for a femur defect mouse model

**DOI:** 10.1101/2020.09.10.289553

**Authors:** Graeme R. Paul, Peter Schwilch, Esther Wehrle, Gisela A. Kuhn, Ralph Müller

## Abstract

Correct fixation of an object is essential for accurate micro computed tomography scanning. In this document, we provide a motivation for, description of, and use cases for a mouse holder appropriate for holding an externally fixated mouse in femur fracture/defect healing experiments. In addition to rigid fixation, the holder provides heating and anaesthetic gas to ensure correct anaesthetic conditions for the animal. We provide the description and design files for a Scanco viva40 scanner, but with small changes, the holder can be used with other scanners.

## Introduction

During animal imaging (in particular microcomputer tomography (micro-CT) imaging) *in vivo*, correct placement and fixation of the animal is essential. Placement is often constrained via the type of scan, the type of scanner and the organ being scanned. While scanning holders are seldom reported in the literature, conventional solutions usually make use of tube-like structures in which the animal’s body part is placed ^1^. However, this often leads to small movements within the scan as well as differing placement positions between scans. To overcome these limitations, we have constructed a mouse holder for a femur defect model that allows rigid, repeatable placement of the organ of interest (the right femur) in the correct position in a micro-CT scan.

The holder provides four core functions:

- Rigid fixation of the femur in the correct location during micro-CT scanning
- Adequate heating of the mouse during scanning
- Delivery of anaesthetic/analgesic gas during scanning
- Interface with additional devices for mechanical intervention experiments

This document provides the manufacturing and use specification of a mouse imaging and loading holder for a femur defect model in mice. The purpose is to hold a mouse, with the right femur fixated with a RISystem external fixator (RISystem, Landquart, Switzerland). This allows repeated measurements of the femur defect and prevents substantial movement between scans, facilitating image registration and longitudinal comparisons^2^. The holder is designed for use with a Scanco viva40 micro-CT scanner (Scanco, Brüttisellen, Switzerland) but with small modifications can be used with other in vivo scanners. Additionally, the holder can be used with a loading machine without the need to transfer the mouse, as the holder can be removed with ease from the scanner.

## Description

**Figure 1:**
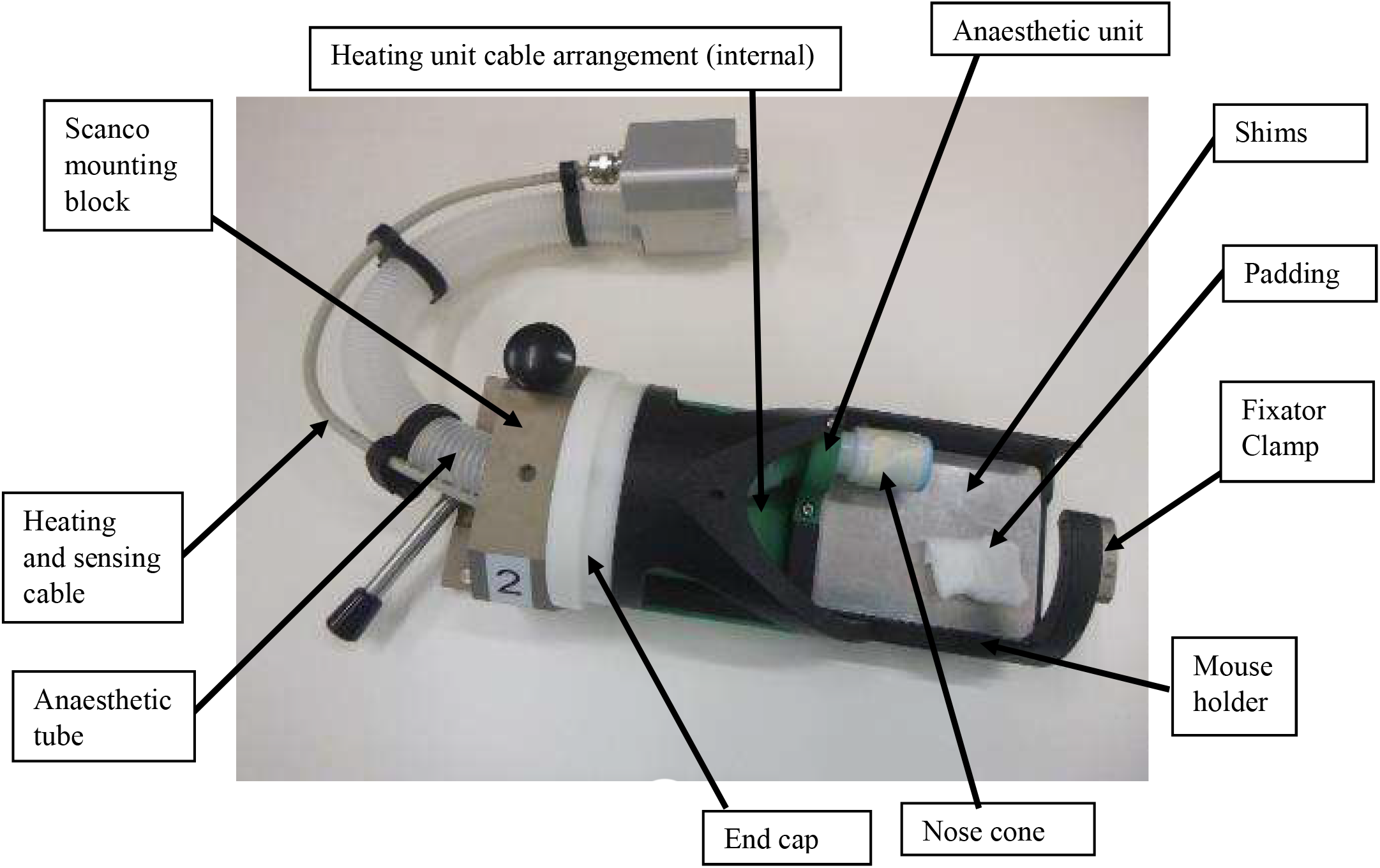
The micro-CT mouse holder

The mouse is placed on the front portion of the heated mouse holder with its nose facing towards the anaesthetic unit (nose cone can be chosen and adjusted according to animal size). The external fixator is placed in a clamp and held via tightening the side screw to a value of 8 Nmm. The fixator and the femur are slightly offset from the centre of the scanning area. Anaesthetic is provided via the anaesthetic unit, consisting of an inner tube for anaesthetic delivery and an outer tube for gas removal. Heating is provided via two 9 W (at 24 V) heating elements of Ø 10 mm flanking a temperature sensor. The unit bolts onto the standard Scanco interface block via the end plate. The full assembly dimensions are 180 mm × Ø 86 mm.

**Figure 2:**
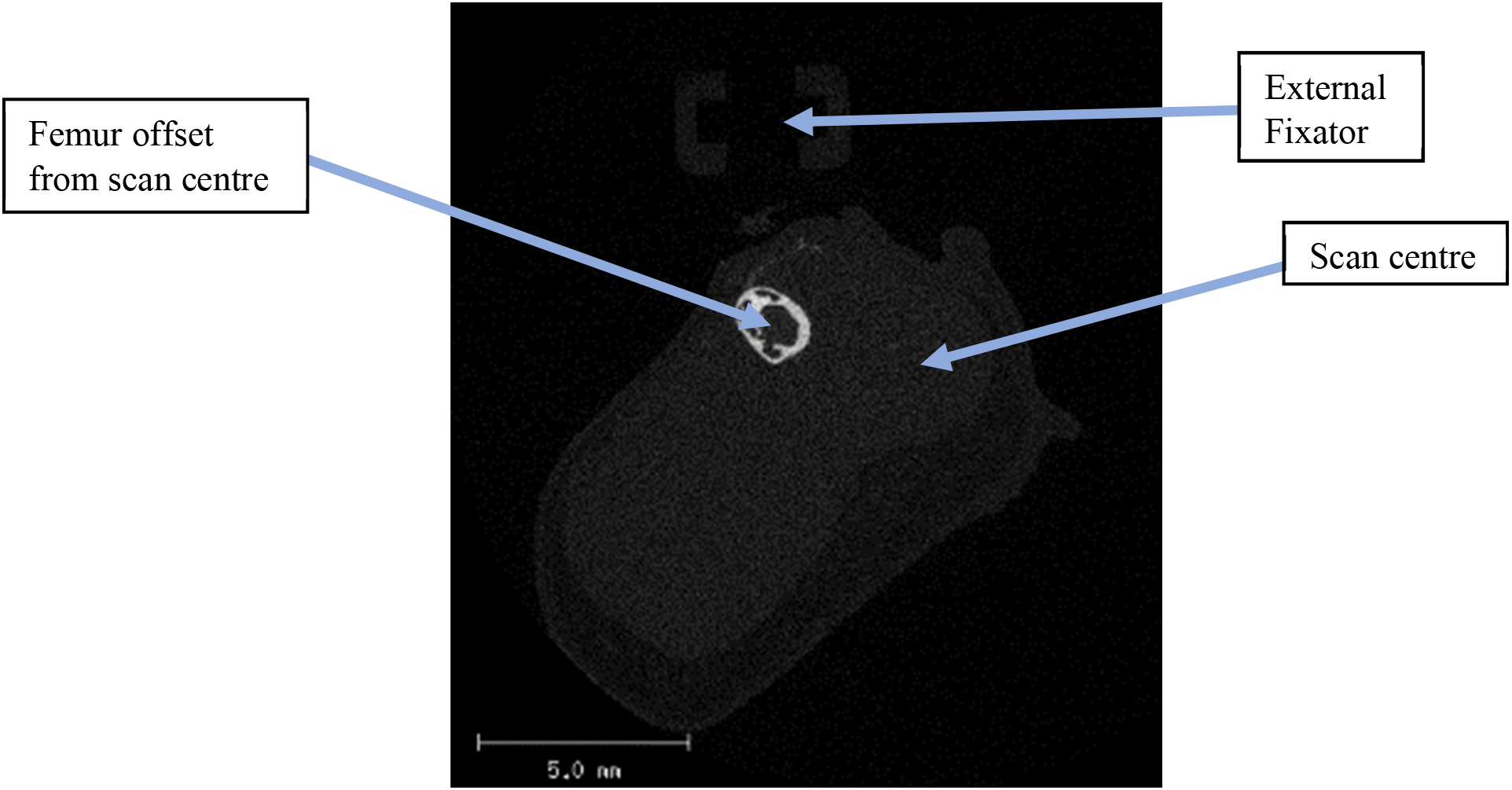
Femur placement in scanned region

### Scanning Location

The femur is positioned off centre and slightly raised above the centre of the scan. The aim of the location is to improve mouse ergonomics, improve scan quality and prevent the muscles and fat from causing local tomography artefacts.

### Part Description

#### Assembly

**Figure 3:**
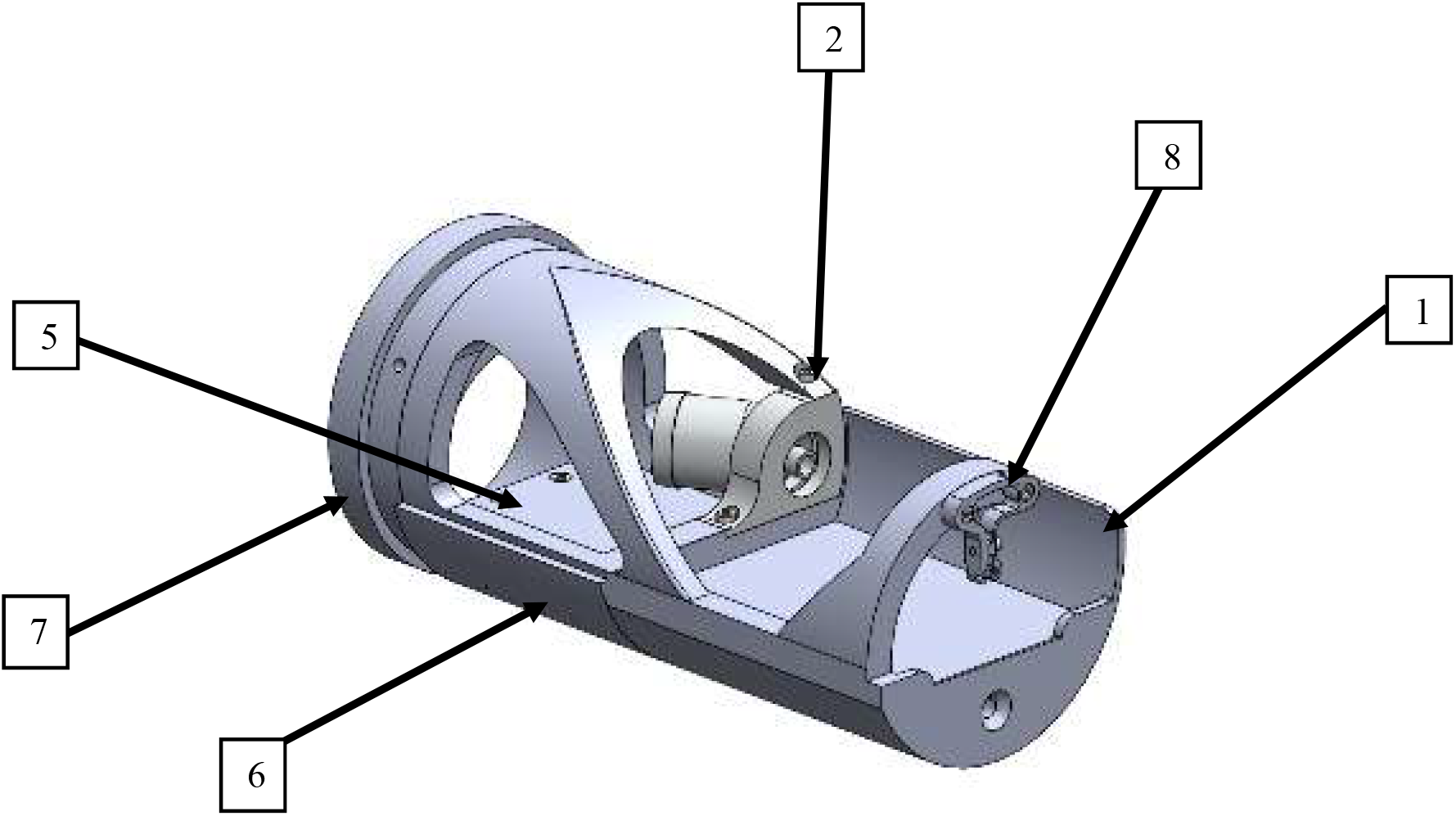
Position of the femur within a micro-CT scan

**Table 1:**
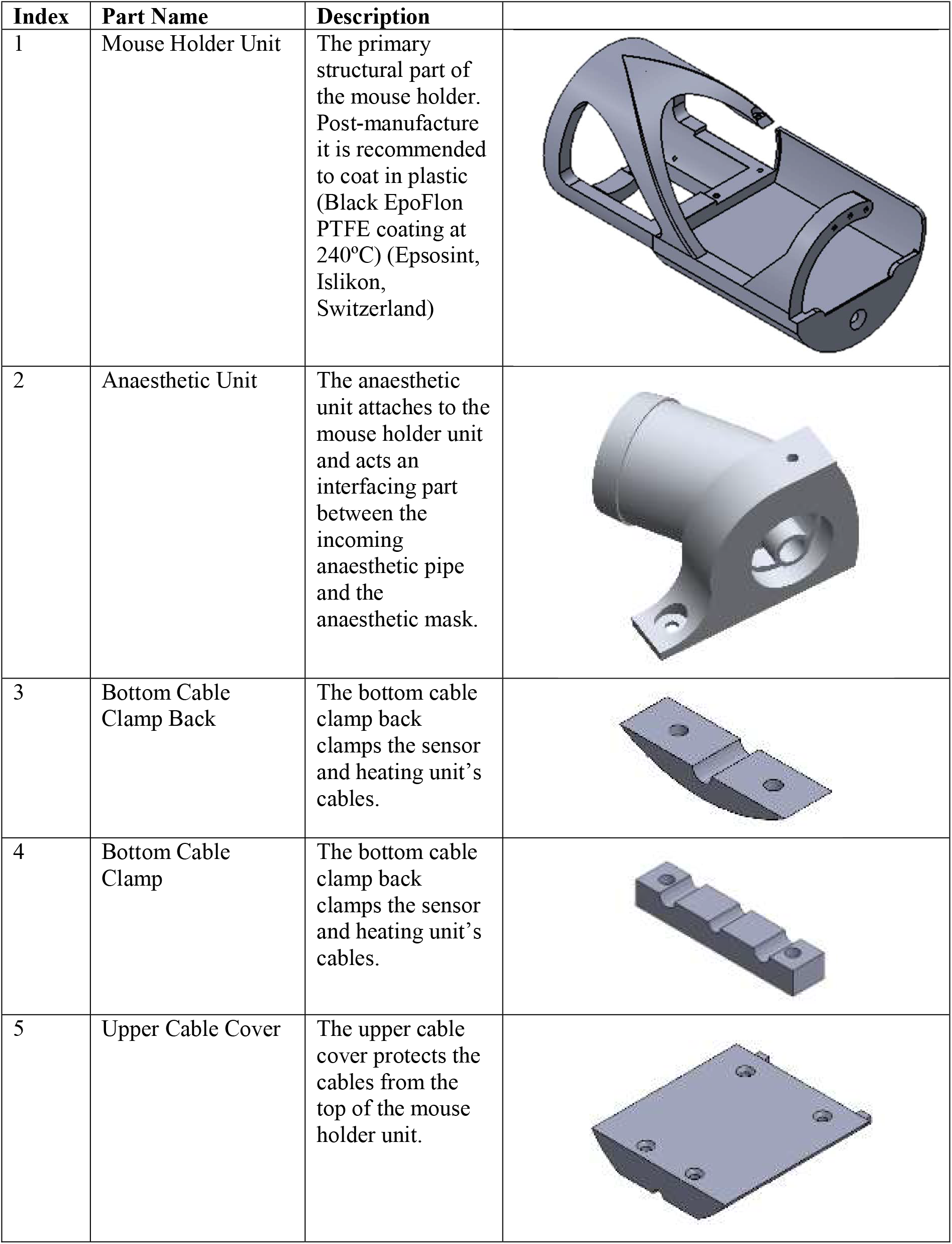
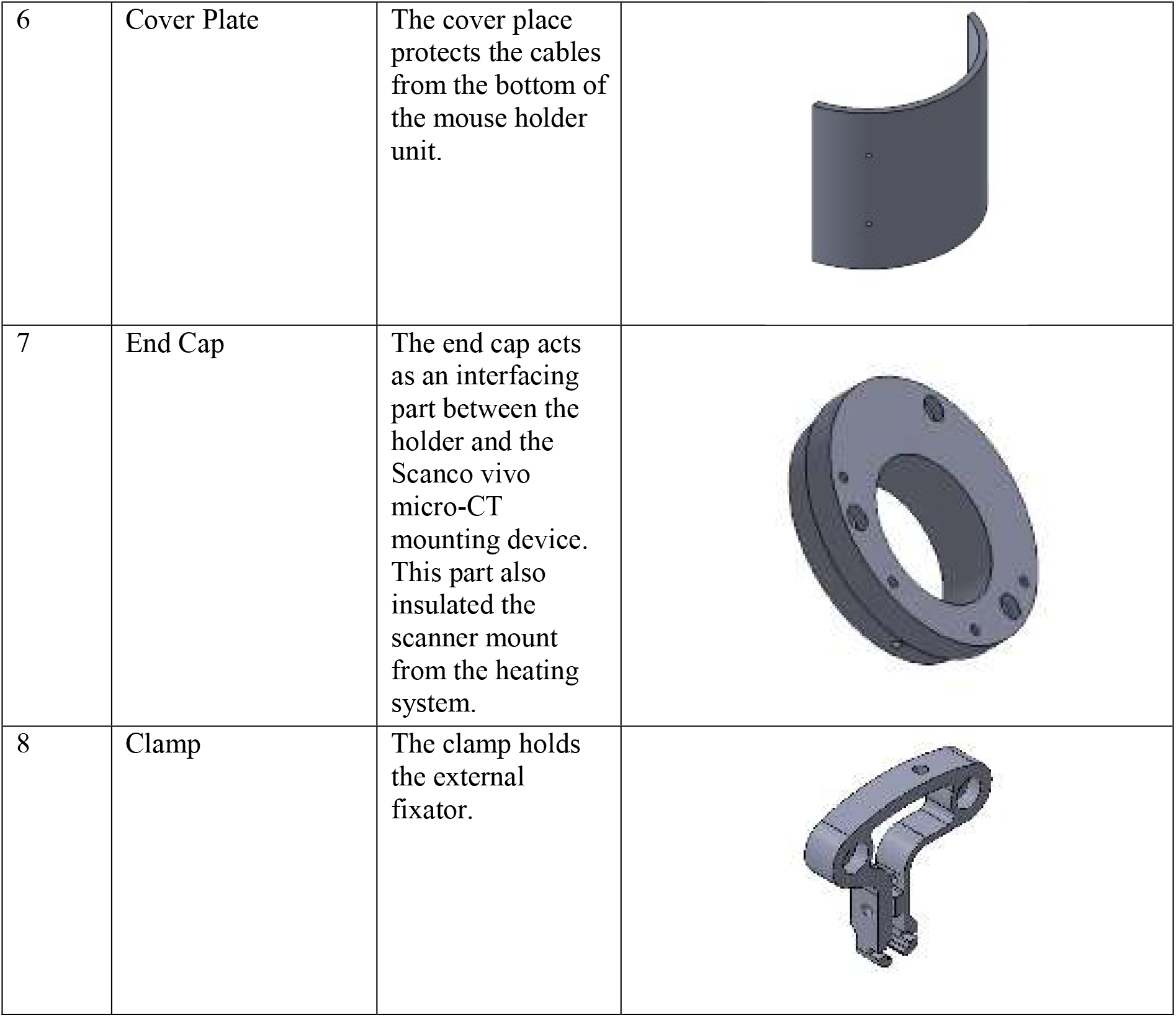
Part numbers, names and description

| Index | Part Name | Description |
| --- | --- | --- |
| 1 | Mouse Holder Unit | The primary structural part of the mouse holder. Post-manufacture it is recommended to coat in plastic (Black EpoFlon PTFE coating at 240°C) (Epsosint, Islikon, Switzerland) |
| 2 | Anaesthetic Unit | The anaesthetic unit attaches to the mouse holder unit and acts as an interfacing part between the incoming anaesthetic pipe and the anaesthetic mask. |
| 3 | Bottom Cable Clamp Back | The bottom cable clamp back clamps the sensor and heating unit's cables. |
| 4 | Bottom Cable Clamp | The bottom cable clamp back clamps the sensor and heating unit's cables. |
| 5 | Upper Cable Cover | The upper cable cover protects the cables from the top of the mouse holder unit. |
| 6 | Cover Plate | The cover place protects the cables from the bottom of the mouse holder unit. |
| 7 | End Cap | The end cap acts as an interfacing part between the holder and the Scanco vivo micro-CT mounting device. This part also insulated the scanner mount from the heating system. |
| 8 | Clamp | The clamp holds the external fixator. |

### Heating and Sensing System

Two Stego PTC Heater RCE 016 (9 W at 24 V) elements are used, flanking a KS-PT1000A-1-550-3-W temperature-sensing unit. These are all controlled via a Jumo CTRON 16 control unit. Both heating elements and the sensor were covered in thermal paste before insertion.

**Figure 4:**
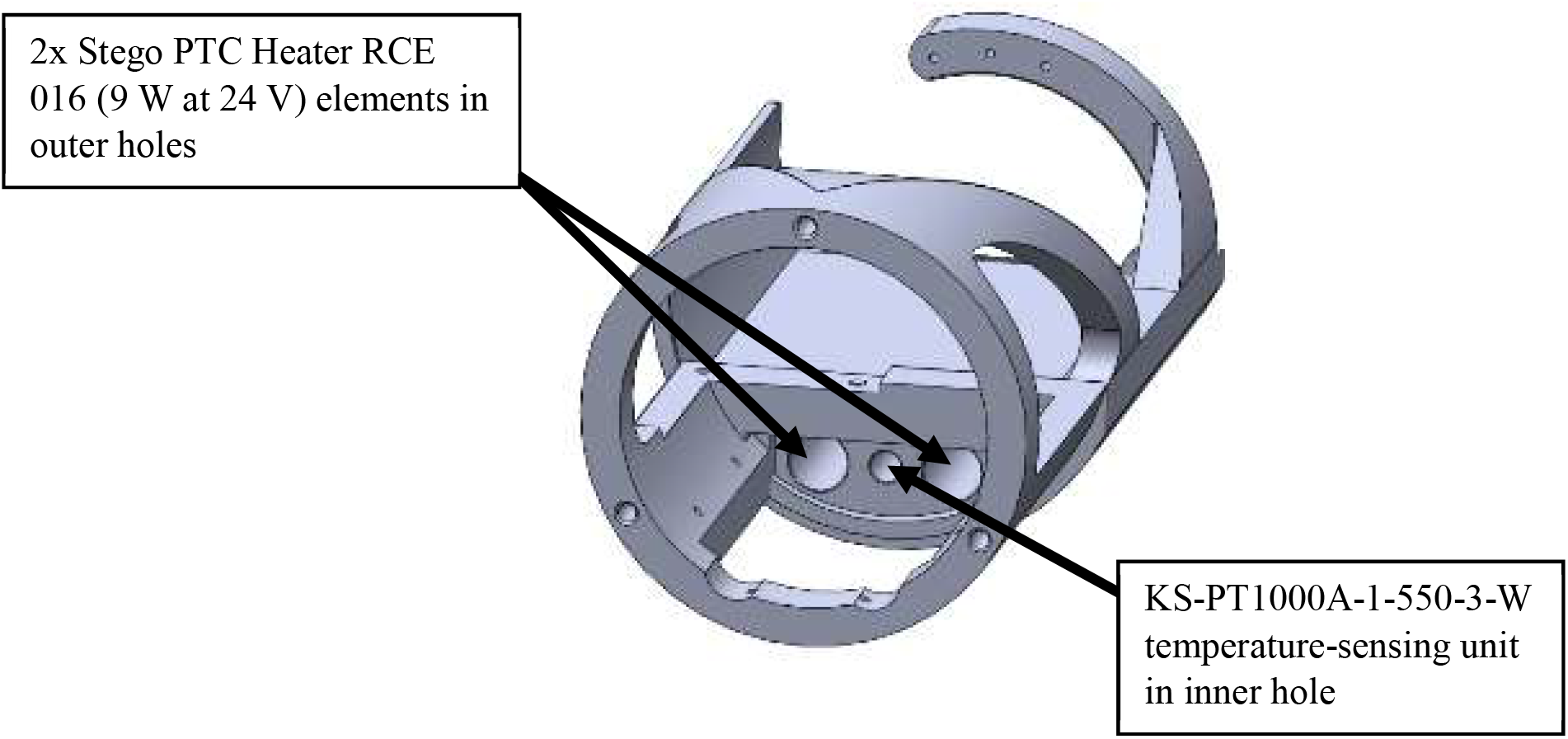
Heating and sensing unit insertion holes

### Examples of use

**Figure 5:**
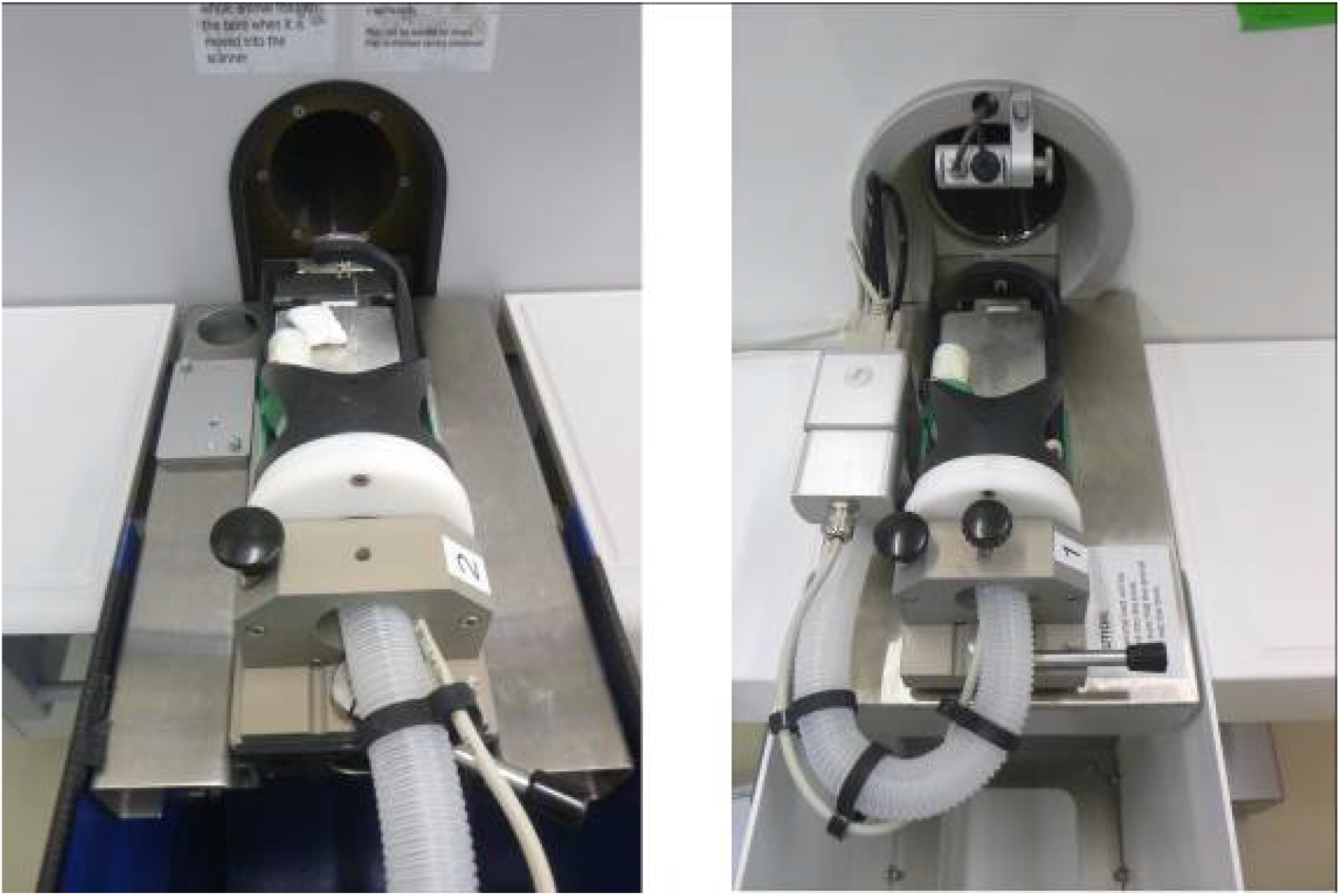
Mouse holder in a Scanco viva40 (left) and a Scanco viva80 (right)

**Figure 6:**
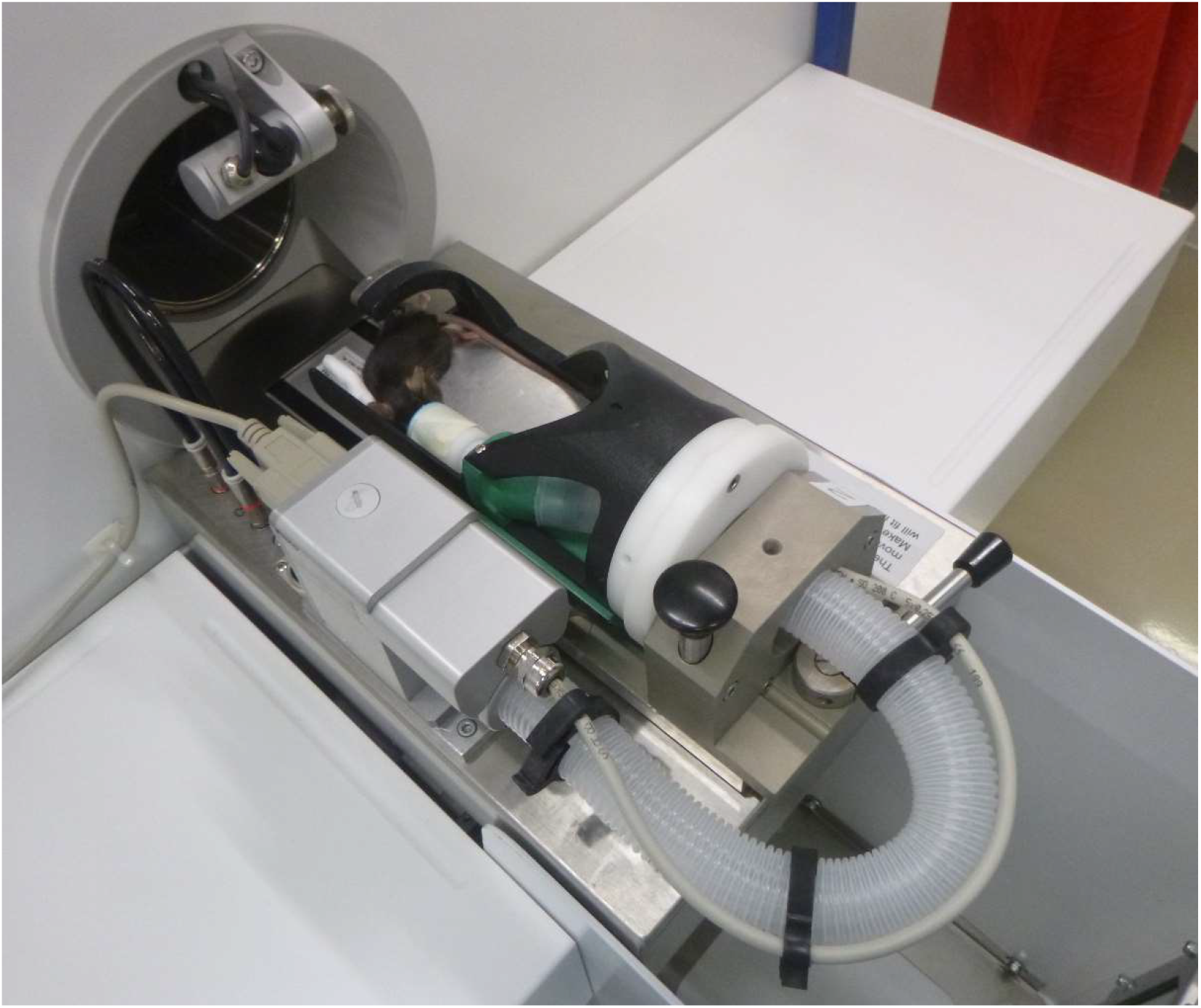
Mouse holder in use in a Scanco viva80 with mouse

**Figure 7:**
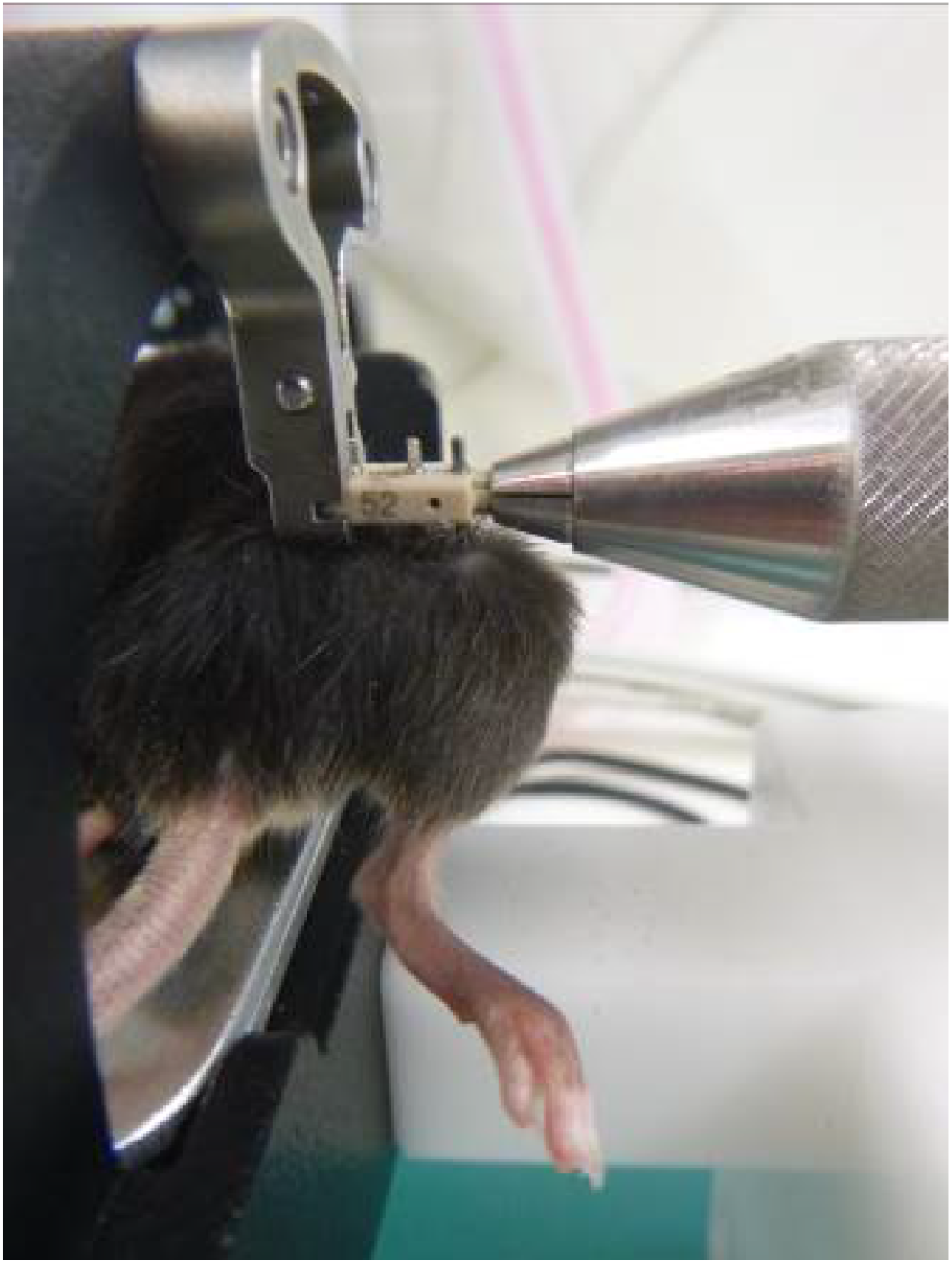
Mouse holder being used in conjunction with a loading device

**Figure 8:**
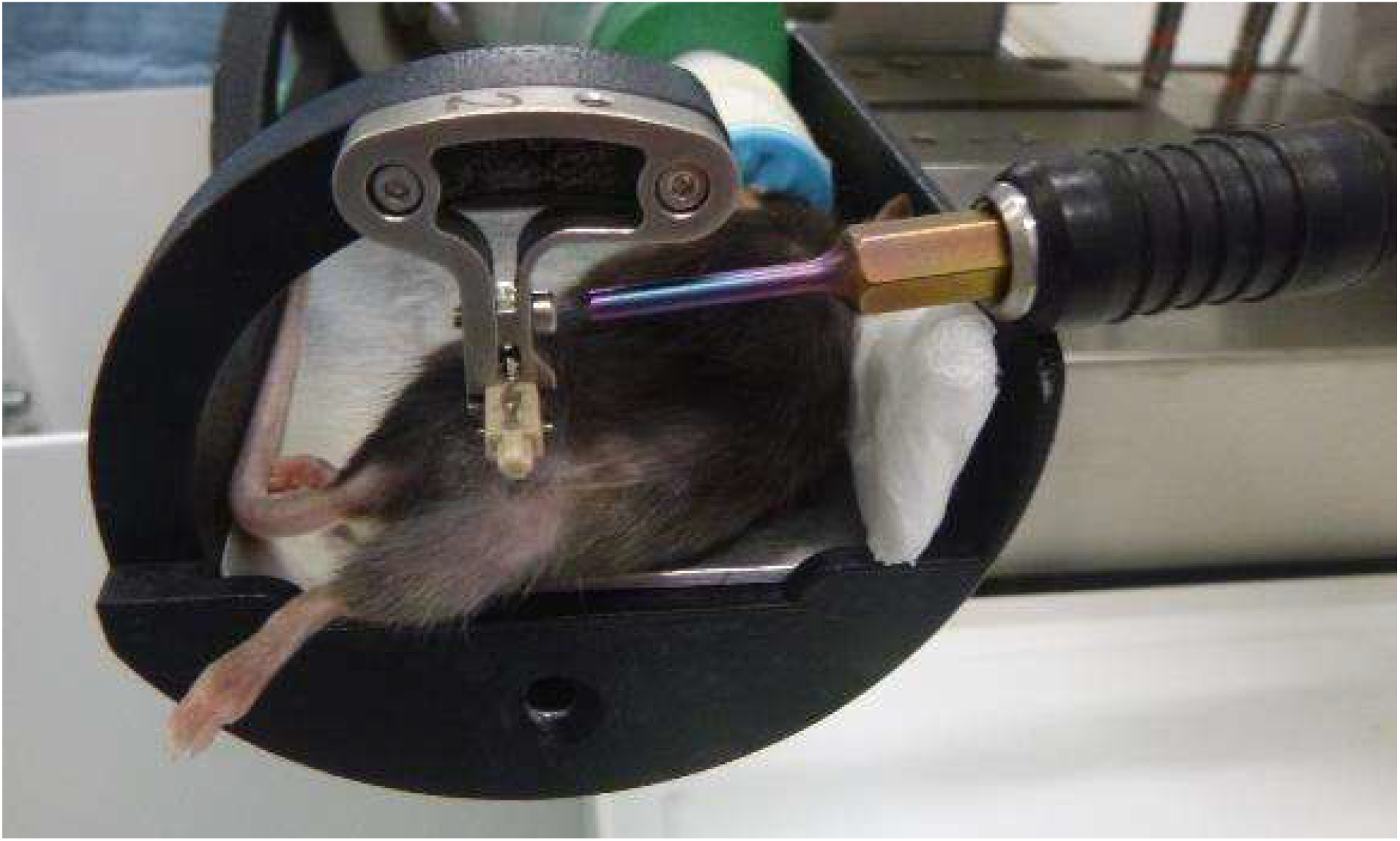
An extended 2.5mm hex torque limited screwdriver is used to secure the external fixator within the clamp. 8 Nmm was used.

**Figure 9:**
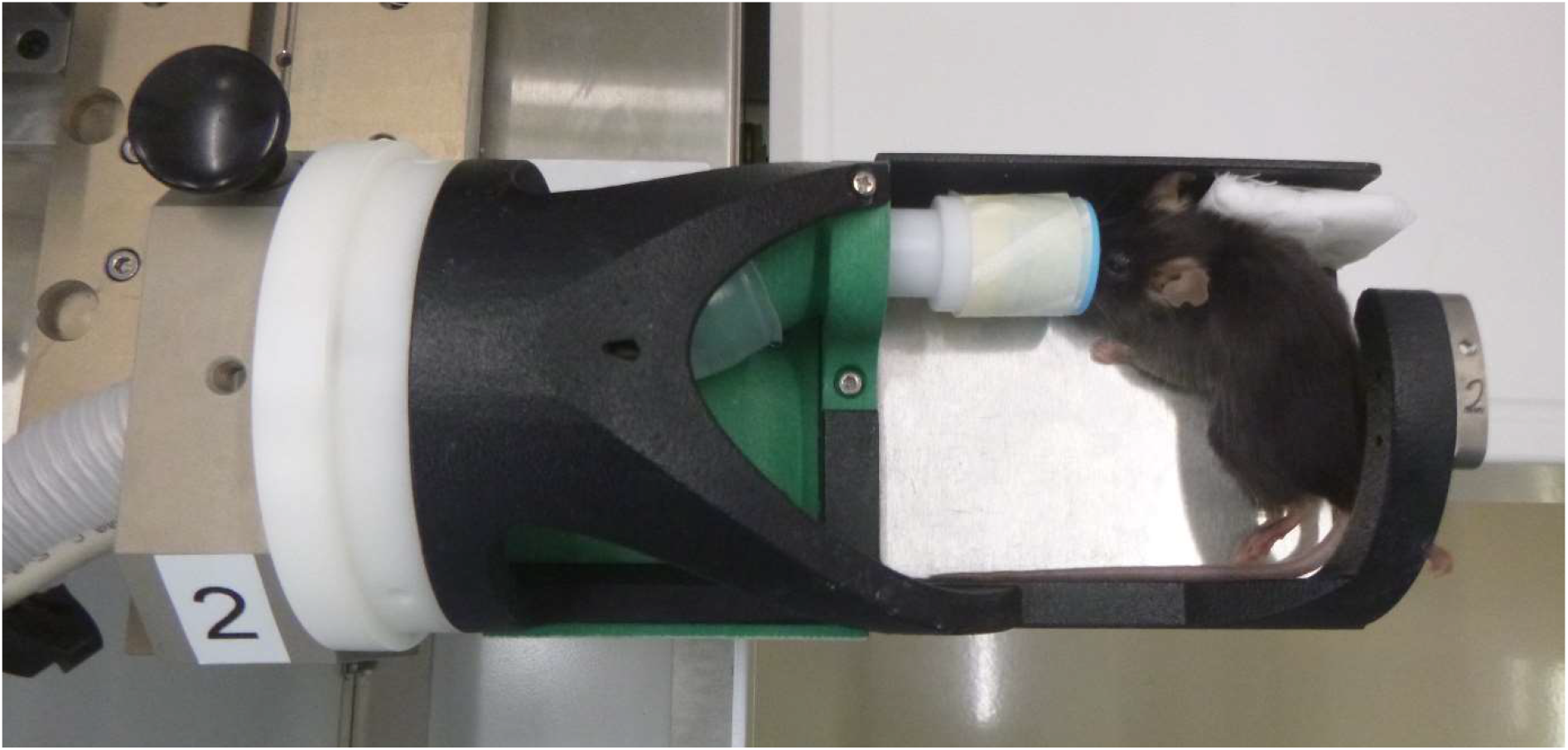
Mouse in mouse holder, top view. Padding is used to improve mouse ergonomics. Shims are used to raise mouse to correct position based on mouse size

## Supporting information

Drawings and Datasheets

## Acknowledgements

We would like to thank Marco Hitz for his assistance on this project. The authors would like to acknowledge support from the European Union (ERC Advanced MechAGE, ERC-2016-ADG-741883). E. Wehrle received funding from the ETH Postdoctoral Fellowship Program (MSCA-COFUND, FEL-25_15-1).

## Appendix A: Mouse holder parts

## Part List

(See accompanying drawings)

| Part | Amount | Material | Manufacturing Process |
| --- | --- | --- | --- |
| Mouse Holder Unit | 1 | Aluminium | Selective Laser Sintering |
| Anaesthetic Unit | 1 | ABS | 3D Printed |
| Bottom Cable Clamp Back | 1 | PMMA | Machined |
| Bottom Cable Clamp | 1 | POM | Machined |
| Upper Cable Cover | 1 | POM | Machined |
| Cover Plate | 1 | POM | Machined |
| End Cap | 1 | ABS | 3D Printed |
| Clamp | 1 | Stainless Steel | Wire cut |
| M2.5x12 | 4 | ISO 7046 | Philips head |
| M2 x 8 | 2 | ISO 4762 | Hex slot head |
| M2.5 x 6 | 1 | ISO 4762 | Hex slot head |
| M4 x 16 | 3 | ISO 4762 | Hex slot head |
| M2 x 6 | 2 | ISO 4762 | Hex slot head |
| Shims (87x67x2mm) | 2 | Aluminium | Plate/Machined |

## Electronic Part List

(See accompanying files)

Electronic Part List (See accompanying files)
| Part | Amount |
| --- | --- |
| Stego PTC Heater RCE 016 (9W at 24 volts) | 2 |
| KS-PT1000A-1-550-3-W | 1 |
| Jumo CTRON 16 control unit | 1 |

## Appendix B: Additional tools

**Figure 10:**
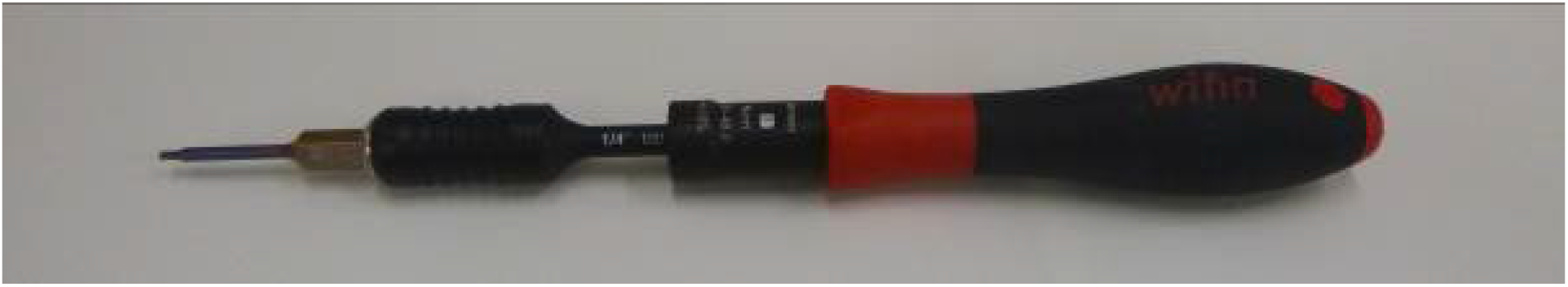
An extended torque limited screwdriver to ensure clamp is not over tightened. 8 Nmm was used.

## Appendix C: Plug and socket

## Description

To facilitate easy transfer of heating and anaesthetic systems a plug and socket system was developed. The socket can be attached to a Scanco viva40 scanner (if the parts labelled **Scanco** are used) or modified appropriately for other scanners or circumstances. The ground plate should be adapted according to the scanning arrangement and scanner mounting point. Depending on the scanner, the parts under **Plug_Socket_Scanco** may need to be used in place of **Plug_Socket**. Scanco appropriate plug-socket parts are prefixed with 1000-05-XXX in their file name. Appropriate mounting plates are included (i.e. Gound_plate_Plug_Socket02-V0.1). All parts were conventionally machined. Part names align with drawing names. Fastener requirements are indicated on the drawings.

## Part photos

**Figure 11:**
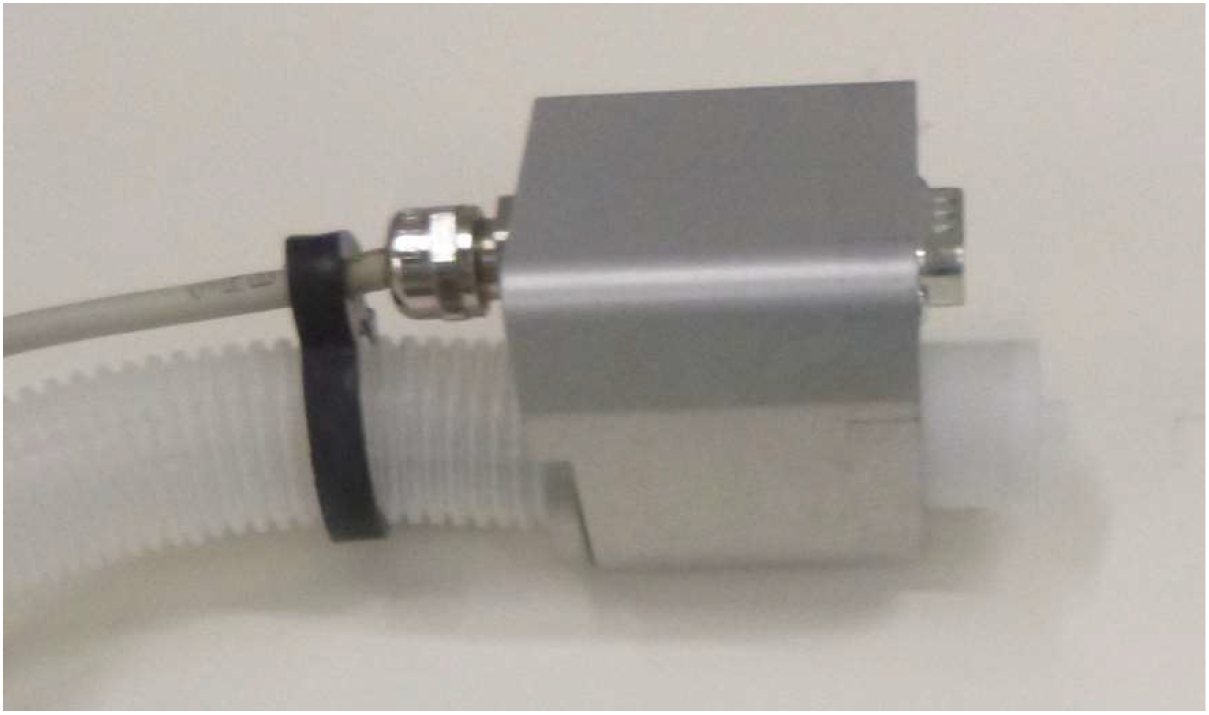
Plug with cables and anaesthetic tube attached.

**Figure 12:**
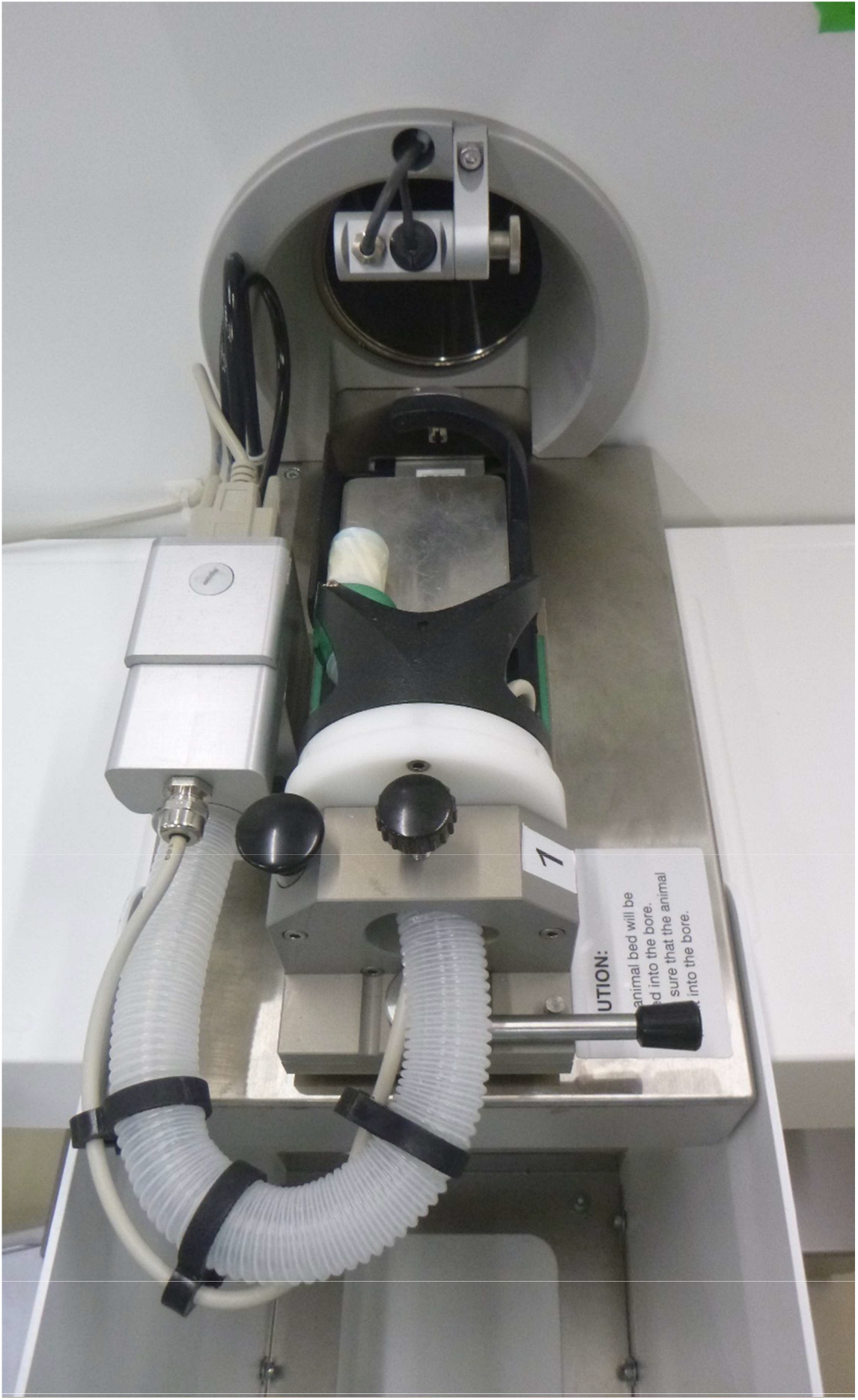
Plug socket arrangement in use

## Part list general

(See accompanying drawings)

| Part | Amount | Material |
| --- | --- | --- |
| Plug | 1 | Aluminium |
| Plug Socket | 1 | Aluminium |
| Plug Socket Cover | 1 | Aluminium |
| Securing cylinder | 1 | Aluminium |
| Anaesthetic mating part | 1 | POM |
| Anaesthetic mating part 2 | 1 | POM |
| 9 pin serial female connector | 1 | - |
| Brass threaded cable joints<br>(M12x1.5) | 1 | - |
| O ring | 1 | - |

## Part list Scanco

| Part | Amount | Material |
| --- | --- | --- |
| Plug-Socket-Scanco | 1 | Aluminium |
| Ground Plate-PlugSocket | 1 | Aluminium |
| Top Plate PlugSocket | 1 | Aluminium |
| PlugSocketCover | 1 | Aluminium |

